# Working memory deficits and altered prefrontal–hippocampal activation in p35 knockout mice with ADHD-like phenotypes: influence of sex and acute monoaminergic treatments

**DOI:** 10.64898/2025.12.23.696253

**Authors:** Florencia Dadam, Osvaldo Martín Basmadjian, Gimena Berardo, Franco Adrián Haehnel, Daiana Yamila Solorzano, María Eugenia Sosa, Sebastián Leonangeli, Andrea Godino, Jorgelina Varayoud, María Gabriela Paglini

**Affiliations:** Laboratory of Neuroplasticity, Instituto de Investigación Médica Mercedes y Martín Ferreyra, INIMEC-CONICET, Universidad Nacional de Córdoba, Córdoba, Argentina; Facultad de Psicología, Universidad Nacional de Córdoba, Córdoba, Argentina; Facultad de Odontología, Universidad Nacional de Córdoba, Córdoba, Argentina; Instituto de Salud y Ambiente del Litoral (ISAL), Consejo Nacional de Investigaciones Científicas y Técnicas (CONICET), Facultad de Bioquímica y Ciencias Biológicas, Universidad Nacional del Litoral (UNL), Ciudad Universitaria, Paraje El Pozo, Santa Fe, 3000, Argentina; Cátedra de Fisiología Humana, Facultad de Bioquímica y Ciencias Biológicas, Universidad Nacional del Litoral (UNL), Ciudad Universitaria, Paraje El Pozo, Santa Fe, 3000, Argentina; Laboratory of Neuroendocrine Regulation of Fluid and Cardiovascular Homeostasis, Instituto de Investigación Médica Mercedes y Martín Ferreyra, INIMEC-CONICET, Universidad Nacional de Córdoba, Córdoba, Argentina; Instituto de Virología ‘’Dr. J. M. Vanella”, Facultad de Ciencias Médicas, Universidad Nacional de Córdoba, Córdoba, Argentina

**Keywords:** CDK5/P35, ANIMAL MODEL, METHYLPHENIDATE, FLUOXETINE, WORKING MEMORY, C-FOS-IR EXPRESSION, SEX EFFECT

## Abstract

Attention-Deficit/Hyperactivity Disorder (ADHD) is a prevalent neurodevelopmental condition characterized by persistent deficits in working memory (WM) and executive control. Dysregulation of the Cyclin-dependent kinase 5 (Cdk5)/p35 signaling pathway has been implicated in ADHD pathophysiology due to its impact on neuronal connectivity and dopamine regulation. Using p35 knockout (p35KO) mice—an animal model exhibiting ADHD-like phenotypes—we investigated WM performance, task-related c-Fos-IR expression under basal conditions as a function of sex and genotype, as well as WM responses to acute treatment with methylphenidate (MPH) or fluoxetine (FLX), administered alone or in combination.

Under basal conditions, p35KO mice exhibited significantly reduced spontaneous alternation in the Y-maze test compared with wild type (WT), whereas recognition memory remained intact. Analysis of c-Fos-IR expression, used as a marker of task-related neuronal activity, revealed region- and genotype-dependent differences, with effects of sex. p35KO animals showed reduced c-Fos-IR expression in prefrontal cortical regions and increased expression in hippocampal regions.

Acute MPH or FLX treatment significantly increased spontaneous alternation in p35KO males, whereas this effect was not observed following combined treatment (MPH+FLX). In WT mice, treatment with MPH, FLX, or MPH+FLX reduced spontaneous alternation, particularly evident in females. Exploratory activity was increased in p35KO mice independently of treatment.

These findings support a role for Cdk5/p35 signaling in the prefrontal-hippocampal activation patterns associated with WM testing and indicate that behavioral and pharmacological responses in this model vary according to sex and neurobiological background, highlighting the importance of considering sex as a biological variable in preclinical and translational ADHD research.

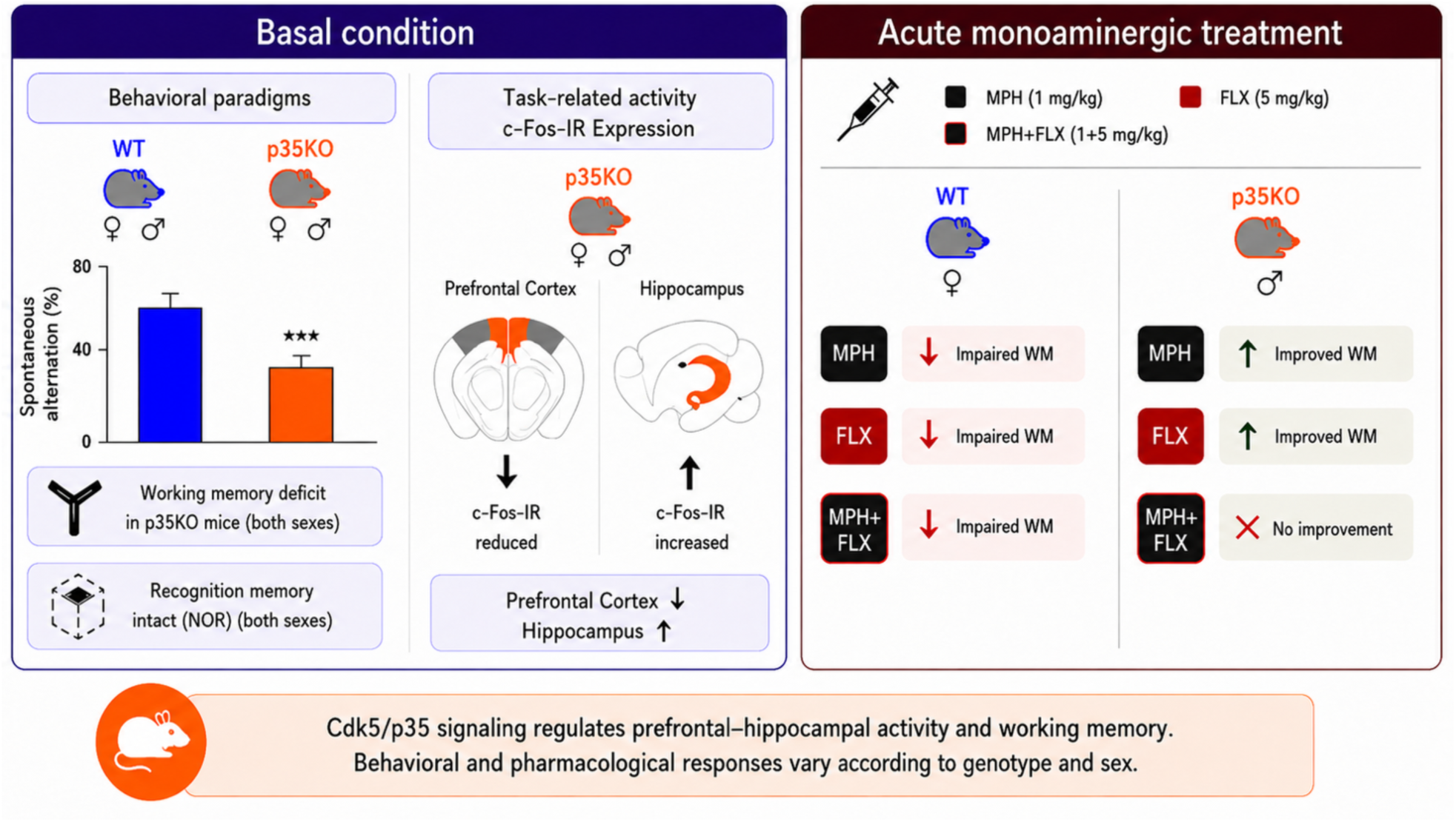

## INTRODUCTION

Attention-Deficit/Hyperactivity Disorder (ADHD) is one of the most prevalent neurodevelopmental conditions in childhood and adolescence, with symptoms often persisting into adulthood. It is characterized by a persistent pattern of inattention, hyperactivity, and impulsivity, which interfere with academic performance, social functioning, and emotional regulation (American Psychiatric Association, 2013).

A deficit in working memory (WM) is one of the most consistently reported cognitive impairments in ADHD, affecting the temporary storage and manipulation of information required for reasoning, planning, and goal-directed behavior (Kofler et al., 2024). WM involves the dynamic interaction of multiple brain regions, particularly the prefrontal cortex (PFC), which orchestrates attentional control and executive functions, and the hippocampus, which contributes to the encoding and retrieval of spatial and contextual information. Altered engagement of prefrontal and hippocampal circuits has been associated with WM deficits observed in ADHD and related disorders (Hou et al., 2022).

The pharmacological treatment of ADHD typically involves the use of psychostimulants, such as methylphenidate (MPH), which enhance dopaminergic and noradrenergic signaling by blocking the reuptake of neurotransmitters. While MPH has proven efficacy in reducing core symptoms of ADHD, its effects are not uniform across individuals, with biological sex emerging as a relevant factor. In cases where ADHD presents with comorbid mood disorders, the selective serotonin reuptake inhibitor (SSRI) fluoxetine (FLX) is sometimes used as an adjunct treatment. However, the cognitive effects of FLX (Chantiluke et al., 2015), alone or in combination with MPH, remain poorly characterized, particularly in developing individuals or animal models representing ADHD-like phenotypes.

A key molecular player in neurodevelopment and cognitive processing is Cyclin-dependent kinase 5 (Cdk5). Unlike other Cdks, Cdk5 requires association with a non-cyclin activator, p35 or p39, to become functionally active (Paglini et al., 1998; Shah and Lahiri, 2017; Svenningsson et al., 2004; Tsai et al., 1994). Alterations in the Cdk5/p35 signaling have been implicated in a variety of neurological and psychiatric conditions (Ferreras et al., 2017; Ikiz and Przedborski, 2008; Kawauchi, 2014; Mlewski et al., 2016, 2008; Rei et al., 2015), including ADHD (Drerup et al., 2010; Fernández et al., 2021; Krapacher et al., 2010), due to its impact on dopamine signaling and neuronal connectivity. Mutant mice lacking p35 (p35 knockout mice, p35KO) represent a preclinical model to investigate ADHD-like phenotypes (de la Peña et al., 2018; Kantak, 2022). Results from our laboratory have shown that these mice exhibit alterations in dopaminergic function, hyperactivity and reduced locomotor activity following MPH or amphetamine treatment—features consistent with core ADHD-related behaviors (Fernández et al., 2021; Krapacher et al., 2010). However, the pharmacological improvement of symptoms, as well as the influence of psychoactive drug type and biological sex, are still poorly understood, despite their potential relevance to treatment efficacy.

Given the interplay between neurotransmitter systems, molecular signaling pathways, and cognitive functions, we investigated whether WM performance and associated prefrontal–hippocampal activation patterns are differentially modulated by acute monoaminergic treatments in the context of ADHD. To this end, we used p35KO mice—a preclinical model exhibiting ADHD-like phenotypes associated with Cdk5/p35 dysregulation that has been primarily characterized in male subjects—to examine cognitive performance and task-related neuronal activation under baseline conditions, as well as the effects of acute drug administration. Furthermore, given that most prior studies using this model have focused on males and the growing evidence of sex-related differences in treatment responses, we included both male and female subjects to better understand biological sources of variability that may contribute to differential therapeutic outcomes in ADHD.

## MATERIAL AND METHODS

### Animals and experimental design in a murine model exhibiting ADHD-like phenotypes

Male and female wild type (WT) and p35 (p35 KO) mutant mice were generated by breeding heterozygous mutants (kind gifts of Dr. L.H. Tsai) (Chae et al., 1997) maintained a C57BL/6J background via brother-sister mating in the vivarium of the INIMEC-CONICET-UNC (Cordoba, Argentina). Animals were weaned at 21 postnatal and housed up to six per cage under a 12-h light/12-h dark cycle and a constant temperature (22° C) with free access to food and water. All the experiments were performed during the light cycle, between 10:00 AM to 3:00 PM in a separated behavioral room. During the test, the room was quiet and slightly lit and mice were allowed to acclimatize for 30 min before the onset of each experiment.

Mice were evaluated during the late postnatal period (PND 22–24), which corresponds to a juvenile developmental stage in rodents characterized by ongoing maturation of cortical and subcortical circuits, particularly within the prefrontal cortex. This developmental window is widely used to model neurodevelopmental processes relevant to the early emergence of cognitive and attentional alterations associated with ADHD (Shaw et al., 2007; Thapar and Cooper, 2016).

All animal procedures and care were approved by the National Department of Animal Care and Health (SENASA – ARGENTINA) and complied with the National Institute of Health guidelines for the Care and Use of Laboratory Animals. All experimental protocols were reviewed and approved by the Institutional Animal Care and Use Committee (CICUAL) at INIMEC-CONICET-UNC. Efforts were made to minimize animal suffering and to reduce the number of animals used.

### Experiment 1: Working memory, exploratory behavior, and c-Fos-IR expression (Y-maze test)

WT and p35KO mice (both males and females) underwent daily handling at postnatal day (PND) 22 to habituate the animals to human contact and minimize stress during subsequent behavioral assessments. On PND 23 mice were evaluated in the Y-maze, a well-established paradigm used to assess spontaneous alternation as a measure of spatial WM and total arm entries as an index of exploratory behavior. Each animal was placed in the maze for 8 min, and spontaneous alternation behavior was recorded as an index of WM performance. 90 min after completing the behavioral test, mice were anesthetized and transcardially perfused with fixative solutions for brain collection (Fig. 1a.). Brains were coronally sectioned and processed for immunohistochemistry of c-Fos immunoreactivity (c-Fos-IR) to quantify region-specific c-Fos-IR expression associated with Y-maze exploration.

**Figure 1.**
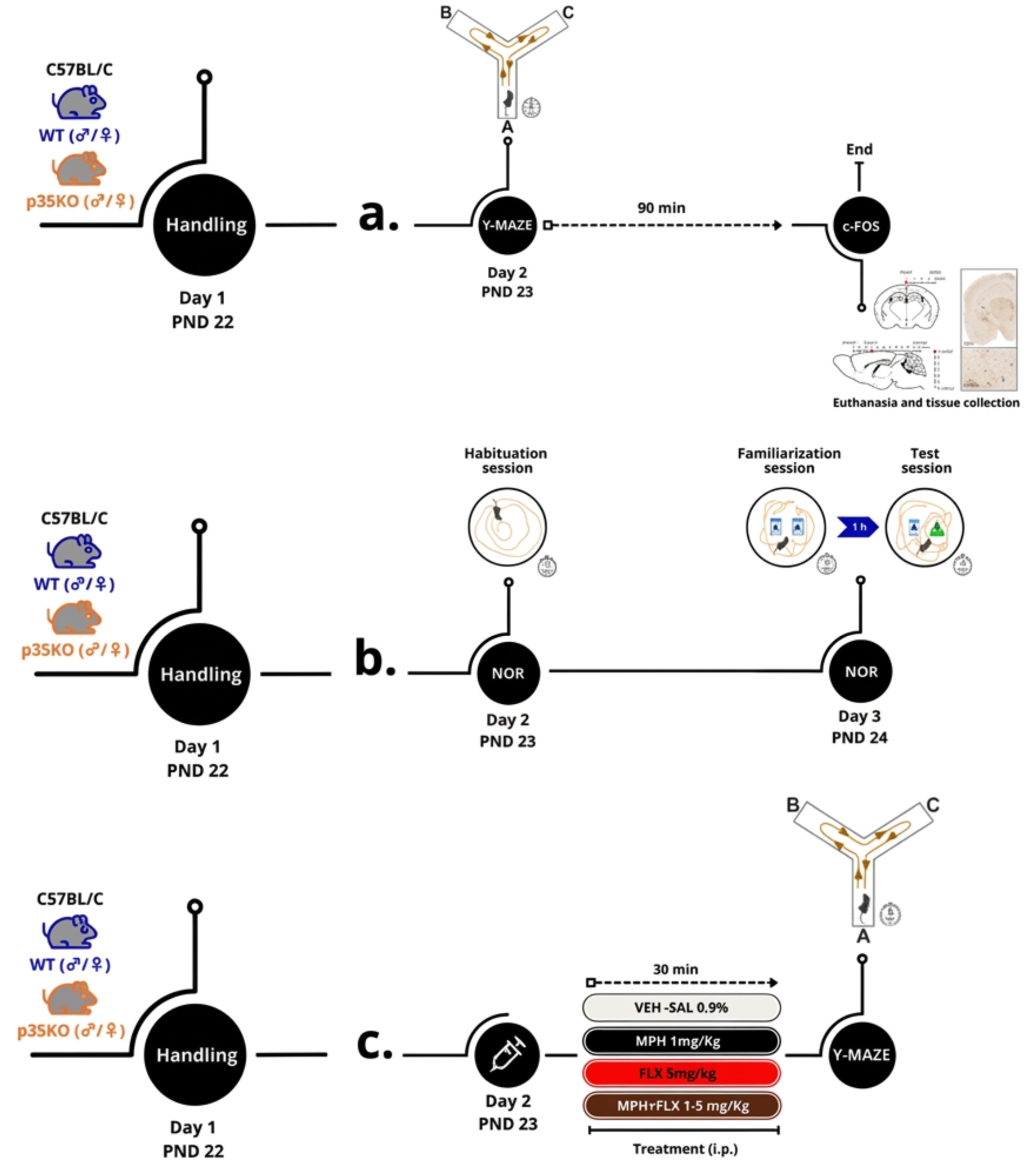
Experimental timelines. (**a) *Y-maze task and brain processing for c-Fos-IR immunohistochemistry.*** WT and p35KO mice (males and females) were handled daily starting at PND 22. At PND 23, animals were tested in the Y-maze for 8 min. 90 min after testing, mice were anesthetized and transcardially perfused for brain collection and preparation of coronal slices for c-Fos-IR immunostaining **(Experiment 1). (b) *Novel Object Recognition (NOR) task.*** WT and p35KO mice (males and females) were handled daily starting at PND 22. On day 2 (PND 23), mice were habituated to the testing arena. On day 3 (PND 24), they underwent the NOR task, consisting of a 10-min familiarization phase with two identical objects (A–A), followed by a 1-h retention interval and a 10-min test phase (A–B). Exploration time of the novel versus the familiar object was quantified as an index of recognition memory **(Experiment 2).** (**c**) ***Y-maze task with psychoactive drug administration***. WT and p35KO mice (males and females) were handled daily starting at PND 22. At PND 23, animals received psychoactive drug treatments and 30 min after drug administration, mice were tested in the Y-maze for 8 min **(Experiment 3).**

### Experiment 2: Recognition memory assessment (Novel Object Recognition Test)

Another group of mice, including WT and p35KO animals of both sexes, received daily handling starting at PND 22 and were subsequently (PND 23) subjected to the Novel Object Recognition (NOR) paradigm, which evaluates recognition memory based on the innate preference of rodents for novelty. On PND 23 (day 2), animals were habituated to the testing arena. On PND 24 (day 3), the task consisted of a 10-min familiarization trial with two identical objects (A–A), followed by a 1-h inter-trial interval. Mice were then reintroduced to the arena for a 10-min test phase in which one familiar object was replaced by a novel object (A–B) (Fig.1b).

### Experiment 3: Psychoactive drugs treatment

Drug treatments were randomly assigned and consisted of a single intraperitoneal (i.p.) injection administered to mice of both sexes (male and female) and genotypes (p35KO and WT) 30 min prior to the Y-maze test (Fig. 1c.). Animals received either methylphenidate (MPH; 1 mg/kg), fluoxetine (FLX; 5 mg/kg), a combination of MPH + FLX (1-5 mg/kg), or Vehicle (VEH; 0.9% saline solution). The MPH used was Ritalin® (Novartis), and FLX was Foxetin® (Gador). The selected doses were based on previous studies. Regarding MPH, this dose increased locomotor activity in WT mice while producing a marked decrease in locomotion in p35KO mice relative to VEH-treated controls (Krapacher et al., 2010). The FLX dose has been shown to alter serotonergic tone and modulate cognitive and emotional behaviors without inducing anxiogenic or sedative effects in rodents (Dhir and Kulkarni, 2007; Hughes, 2004; Popa et al., 2010). All procedures involving substance administration were approved by resolution 021-2017B of the Institutional Committee for the Care and Use of Laboratory Animals (CICUAL) at the Instituto de Investigación Médica Mercedes y Martín Ferreyra.

### Behavioral tests

#### Y-maze test (Y-MAZE)

The Y-maze apparatus was constructed of white-painted plywood, with three arms (50 cm long × 4 cm wide × 12 cm high) positioned at 120° angles. Each mouse was placed at the end of one arm facing the center and allowed to explore freely for 8 min. The maze was cleaned with 70% ethanol between trials to eliminate olfactory cues.

Arm entries were defined as the crossing of all four paws from the central area into a given arm, with the animal’s snout oriented toward the distal end. Spontaneous alternation behavior was defined as consecutive entries into the three different arms in overlapping triplet sets. The percentage of spontaneous alternation was calculated as the ratio of actual to possible alternations (total arm entries − 2), multiplied by 100, as previously described (Hughes, 2004; Miedel et al., 2017). Total arm entries were used as an index of exploratory activity. All behavioral recording and scoring were performed manually by the same experimenter, blinded to genotype and treatment conditions, to ensure procedural consistency and standardization.

#### Novel object recognition task (NOR)

The NOR test was performed to evaluate short-term recognition memory based on spontaneous exploratory behavior. The task consisted of three phases: habituation (10 min free exploration of an empty arena), familiarization/acquisition (10 min exploration of two identical objects), and test/retention (1 h later, one familiar object replaced with a novel one). The objects differed in shape but were otherwise identical in size, material, color, and odor, and were not climbable. Trials were conducted in a circular arena (40 cm diameter under dim light conditions (∼40 lx). Object identity and location were counterbalanced across animals to avoid spatial or object bias. Exploration was operationally defined as the period during which the animal’s nose remained within ≤ 2–3 cm of the object accompanied by active vibrissae movement, consistent with established criteria. The time spent exploring the novel and familiar object was quantified during both the familiarization and test sessions using an automated tracking script implemented in Fiji (ImageJ), ensuring objective and standardized measurement. Recognition index was calculated as: (time exploring the novel object / total exploration time) × 100, where total exploration time corresponds to the sum of time spent exploring both objects. Animals with total exploration time < 20 s during the test session were excluded from analysis. The automated Fiji/ImageJ-based tracking script was validated against manual scoring in a subset of sessions.

### c-Fos-IR immunohistochemistry and quantification

c-Fos-IR expression was used as a marker of task-related neuronal activity and was assessed by immunohistochemistry following established procedures (Dadam et al., 2014). WT and p35KO mice were anesthetized with 30% chloral hydrate and transcardially perfused with saline followed by 4% paraformaldehyde. Brains were post-fixed, cryoprotected, and cut into 40-µm coronal sections. Sections were incubated with a rabbit anti–c-Fos antibody (Ab-5, Oncogene Science) and processed using a biotinylated secondary antibody, avidin–biotin complex, and DAB for visualization. Brain section images were acquired using a PhenoImager™ Fusion microscope (Akoya Biosciences) and visualized with Phenochart software (v1.2.0), which incorporates multirange spectral detection and multispectral imaging (MSI) technology. Quantification was performed using a custom ImageJ macro that automated image preprocessing, ROI alignment, and particle detection. c-Fos-IR nuclei were quantified in cortical (+2.22 mm) and hippocampal (−1.58 mm) regions across four predefined anteroposterior levels relative to Bregma, according to stereotaxic coordinates(Paxinos and Franklin, 2019). The macro generated standardized outputs for nuclei count, nuclear area, and ROI size. Full details of anatomical ROI definitions, preprocessing steps, image-processing parameters, and the complete macro code are provided in the Supplementary Material and Methods.

### Statistical analysis

Statistical analyses were performed using Statistica Stat-Soft Inc. version 8.0 Tulsa, OK, USA. Assumptions of homoscedasticity and normality were tested using Levene and Shapiro-Wilks tests, respectively. Data were analyzed using two-way ANOVA (sex × genotype) or three-way ANOVA (sex × genotype × treatment), as appropriate, followed by Tukey’s post hoc tests. Main effects and interactions were evaluated for each dataset. In cases where no significant interactions involving sex were detected, results were interpreted based on main effects. Statistical significance was set at p < 0.05. Data are presented as the mean ± standard error of the mean (SEM).

## RESULTS

### Impairment of Working Memory in Mice Lacking Cdk5-activator p35

Given the relevance of WM deficits in ADHD, we first investigated this cognitive domain in WT and p35KO mice under basal conditions. To this end, sex- and genotype-related differences in cognitive performance were analyzed in the spontaneous Y-maze test using a two-way ANOVA with sex and genotype as factors.

A significant main effect of genotype was observed (F(1, 21) = 32.04, p < 0.0001), with p35KO mice showing reduced alternation compared to WT animals. Post hoc comparisons revealed that both male and female p35KO mice exhibited WM impairment, as evidenced by a reduction in the percentage of alternation compared to their respective WT controls. No significant sex × genotype interaction was detected.

In addition, a significant main effect of sex was observed (F(1, 21) = 11.65, p < 0.049), independent of genotype, with females showing a higher percentage of alternation than males (Fig. 2a). To assess exploratory activity, the total number of arm entries was analyzed. A significant main effect of genotype was observed (F(1, 21) = 23.55 p < 0.001), with p35KO mice displaying a higher number of arm entries compared to WT animals, consistent with increased exploratory activity in this animal model (Fig. 2b).

**Figure 2.**
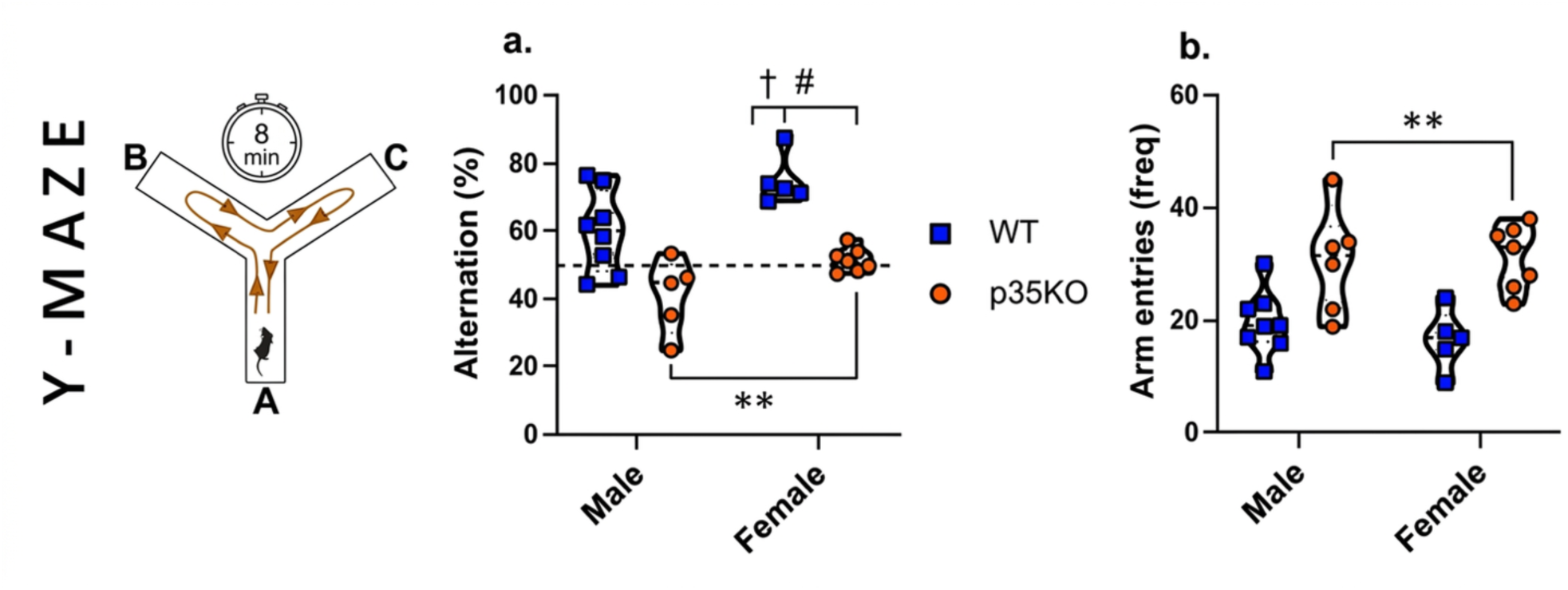
Cognitive and exploratory performance of WT and p35KO mice of both sexes in the Y-maze task. **Alternation percentage (a):** Both male and female p35KO mice displayed decreased alternation performance compared with their respective WT controls, consistent with reduced WM performance (n=6-7, **p < 0.01). A significant main effect of sex was also observed, with females exhibiting higher percentage of alternation than males, regardless of genotype (#p < 0.05). **Arm entry frequency (b):** Both male and female p35KO mice exhibited an increased number of arm entries relative to WT mice, consistent with enhanced exploratory activity (n=6-7, **p < 0.01). Data are expressed as mean ± SEM. Statistical analyses were performed using two-way ANOVA followed by Tukey’s post hoc test.

### Sex-Related Differences in Exploratory Behavior with Preserved Recognition Memory in p35 KO and WT Mice

To assess hippocampal-dependent aspects of recognition memory in p35KO and WT mice, we employed the Novel Object Recognition (NOR) test, which capitalizes on rodents’ innate preference for novelty. This task is widely used to evaluate responses to novel stimuli and the ability to discriminate between familiar and unfamiliar objects (Broadbent et al., 2004; Ennaceur and Delacour, 1988).

Exploratory behavior during the training and test sessions was analyzed using a two-way ANOVA, with sex and genotype as factors. In both the training and test sessions, the recognition index was significantly above chance level (0.5) in all groups, with no main effects of genotype or sex, and no interaction between factors. These results suggest that recognition memory is preserved independently of genotype and sex (Fig. 3a and c).

**Figure 3:**
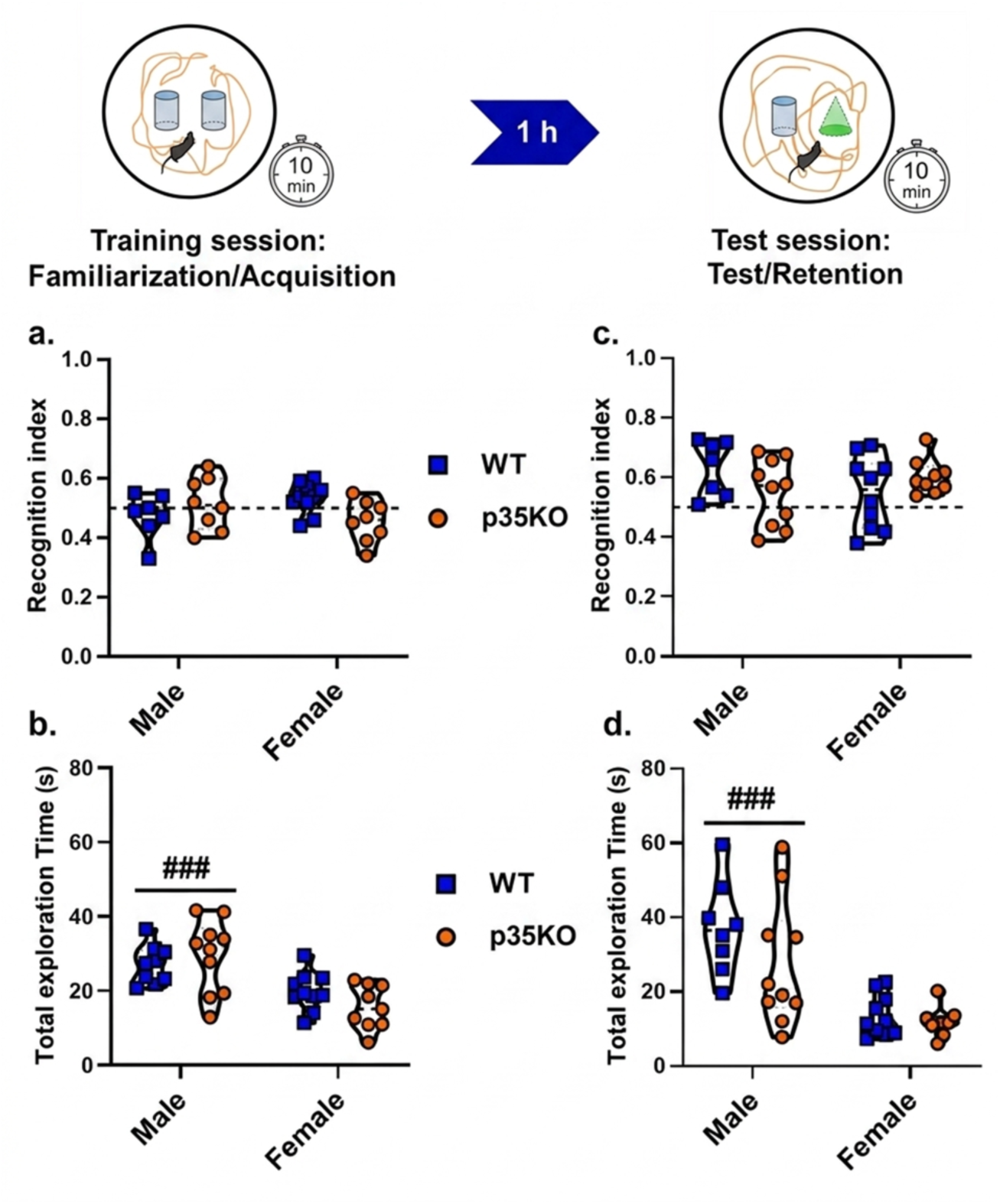
Behavioral performance of WT and p35KO mice of both sexes in the Novel Object Recognition (NOR) task. Recognition index for the novel object during the training (a) and test (c) sessions across experimental groups: No significant effects of genotype or sex were observed. Total exploration time in seconds (s) during the training (b) and test (d) sessions across experimental groups. A significant main effect of sex was found both in the training session (n=16-17, ###p = 0.001) and in the test session (n=16-17, ###p = 0.0001). Data are expressed as mean ± SEM for WT males, WT females, p35KO males, and p35KO females.

Analysis of total exploration time in both sessions revealed a significant main effect of sex on total exploration time [training session: F(1, 62) = 15.62, p = 0.001; test session: F(1, 62) = 25.71, p < 0.0001], with females—regardless of genotype—spending less time exploring the objects compared to males (Fig. 3b and d).

Taken together, these findings suggest that although total exploratory behavior is modulated by sex, recognition memory—as assessed by the NOR task—is not significantly altered by the absence of p35.

### Differential prefrontal and hippocampal c-Fos-IR expression patterns in p35KO mice following working memory testing

Task-related neuronal activity was inferred from c-Fos-IR expression, which was quantified in multiple prefrontal and hippocampus regions, 90 min after the Y-maze test (Fig. 4 and S1, S2). The quantification of the number of c-Fos-IR positive cells was performed in CA1, CA3 and dentate gyrus of hippocampus (Fig 4 a, b, c and S1) and prelimbic (PL), infralimbic (IL), Lateral Orbitofrontal cortex (LO), Cingulate (Cg1) and Ventro orbital (VLO) of prefrontal cortex (Fig 4 d, e, f, g, h and S2).

**Figure 4.**
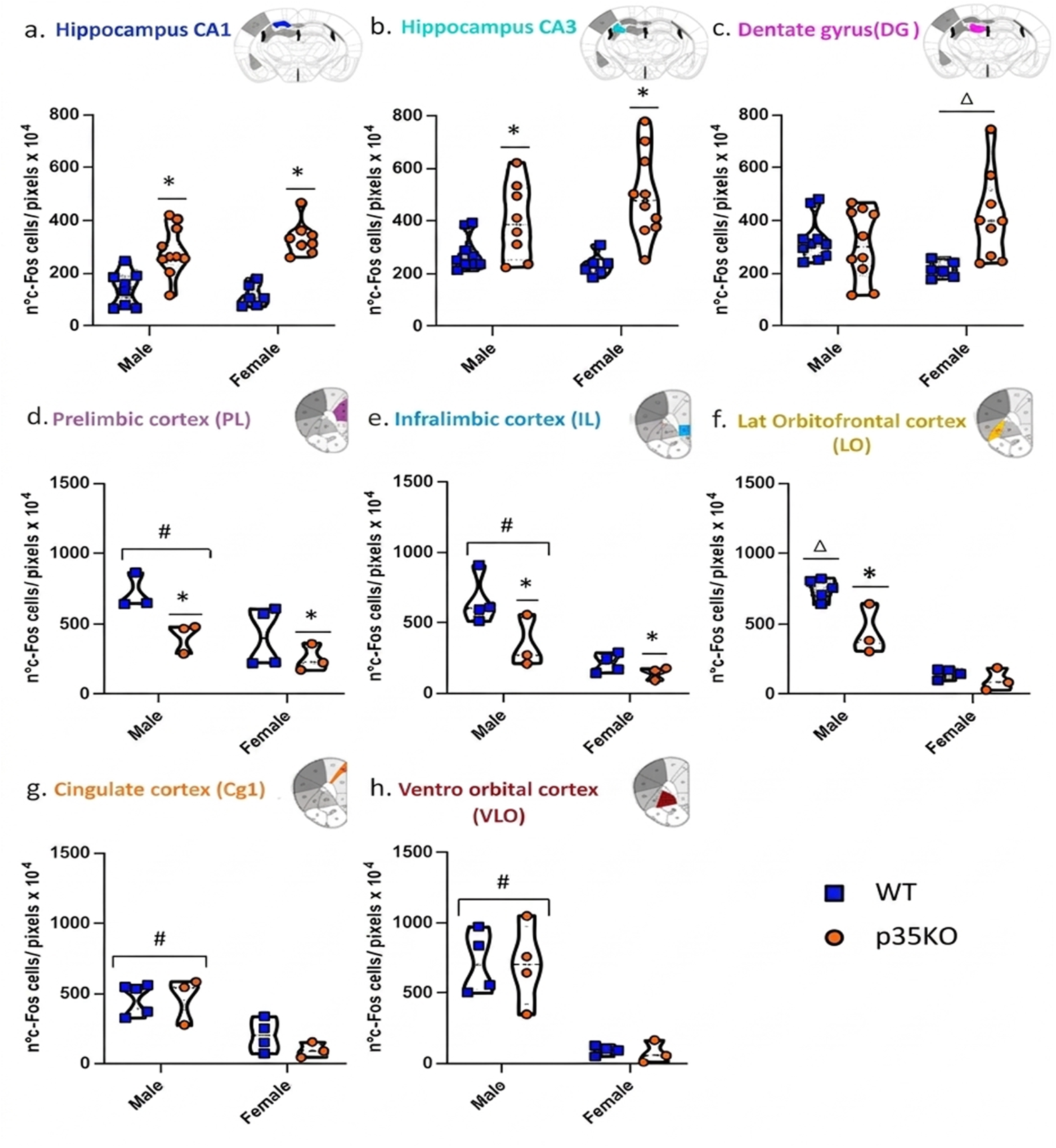
Differences in c-Fos-IR expression across hippocampal and prefrontal cortex subregions in WT and p35KO mice of both sexes. a) **CA1:** *p < 0.01, main effect of genotype (WT vs. p35KO). (b) **CA3:** *p < 0.01, main effect of genotype (WT vs. p35KO). (c) **Dentate gyrus (DG):** Δp < 0.01, p35KO females vs. WT females. (d) **Prelimbic cortex (PL):** #p < 0.05, main effect of sex (males vs. females); *p < 0.05, main effect of genotype (WT vs. p35KO). (e) **Infralimbic cortex (IL):** #p < 0.01, main effect of sex (males vs. females); *p < 0.05, main effect of genotype (WT vs. p35KO). (f) **Lateral orbitofrontal cortex (LO):** Δp < 0.05, WT males vs. all other groups; ¥ p < 0.05, p35KO males vs. p35KO females. (g) **Cingulate cortex (Cg1):** #p < 0.01, main effect of sex (males vs. females). (h) **Ventrolateral orbitofrontal cortex (VLO):** #p < 0.01, main effect of sex (males vs. females). Data are expressed as mean ± SEM. Statistical analyses were performed using two-way ANOVA followed by Tukey’s post hoc test.

The expression of c-Fos-IR was significantly reduced in p35KO animals across several prefrontal subregions [PL: F(1, 9) = 7.363, p < 0.05; IL: F(1, 10) = 6.838, p < 0.05), consistent with lower neuronal activation in this genotype. In addition, a significant main effect of sex was observed [PL:(F(1, 9) = 7.802, p < 0.05; IL: F(1, 10) =20.32, p < 0.01: VLO: F(1, 12) = 19.03, p < 0.01; Cg1: F(1, 11) = 26.54, p < 0.01] with males exhibiting higher c-Fos-IR expression than females independently of genotype across most of the prefrontal cortex regions analyzed.

Post hoc comparisons revealed region-specific differences. In the (LO), WT males showed higher c-Fos-IR expression compared to all other groups, and p35KO males exhibited higher levels than p35KO females (F(1, 11) = 6.251, p < 0.05). These findings indicate a progressive reduction in c-Fos-IR expression across groups, with the lowest levels observed in p35KO females (Fig. 4d–h and S2).

Conversely, analysis of c-Fos-IR expression in the hippocampus revealed that both male and female p35KO mice exhibited significantly higher expression in the CA1 and CA3 regions compared with WT counterparts (CA1: F(1, 28) = 48.17, p < 0.01; CA3: F(1, 30) = 19.93, p < 0.01), consistent with increased neuronal activation in these areas of the dorsal hippocampus (Fig 4 a and b). Moreover, a significant increase in c-Fos-IR expression was observed in the DG exclusively in female p35KO compared with WT female mice (F(1, 31) = 6.847, p < 0.05) (Fig. 4 c). Taken together, these results indicate region-specific differences in c-Fos-IR patterns between prefrontal and hippocampal regions in p35KO mice following WM testing.

### Opposite effects of psychoactive drugs on working memory performance in p35KO and WT mice

Working memory was assessed in the Y-maze 30 min after psychoactive drugs administration (MPH, FLX, or MPH+FLX). Performance was analyzed using a three-way ANOVA including sex, genotype, and treatment as factors. No significant interactions involving sex were detected; therefore, data are displayed separately by sex for visualization purposes only. The analysis revealed a significant genotype × treatment interaction (F(3,54)=3.72, p=0.0166).

Under vehicle-treated conditions, p35KO mice of both sexes (p < 0.001) exhibited reduced spontaneous alternation compared to WT controls, consistent with the deficit observed in Experiment 1 and replicated across cohorts (Fig. 5 a, c).

**Figure 5.**
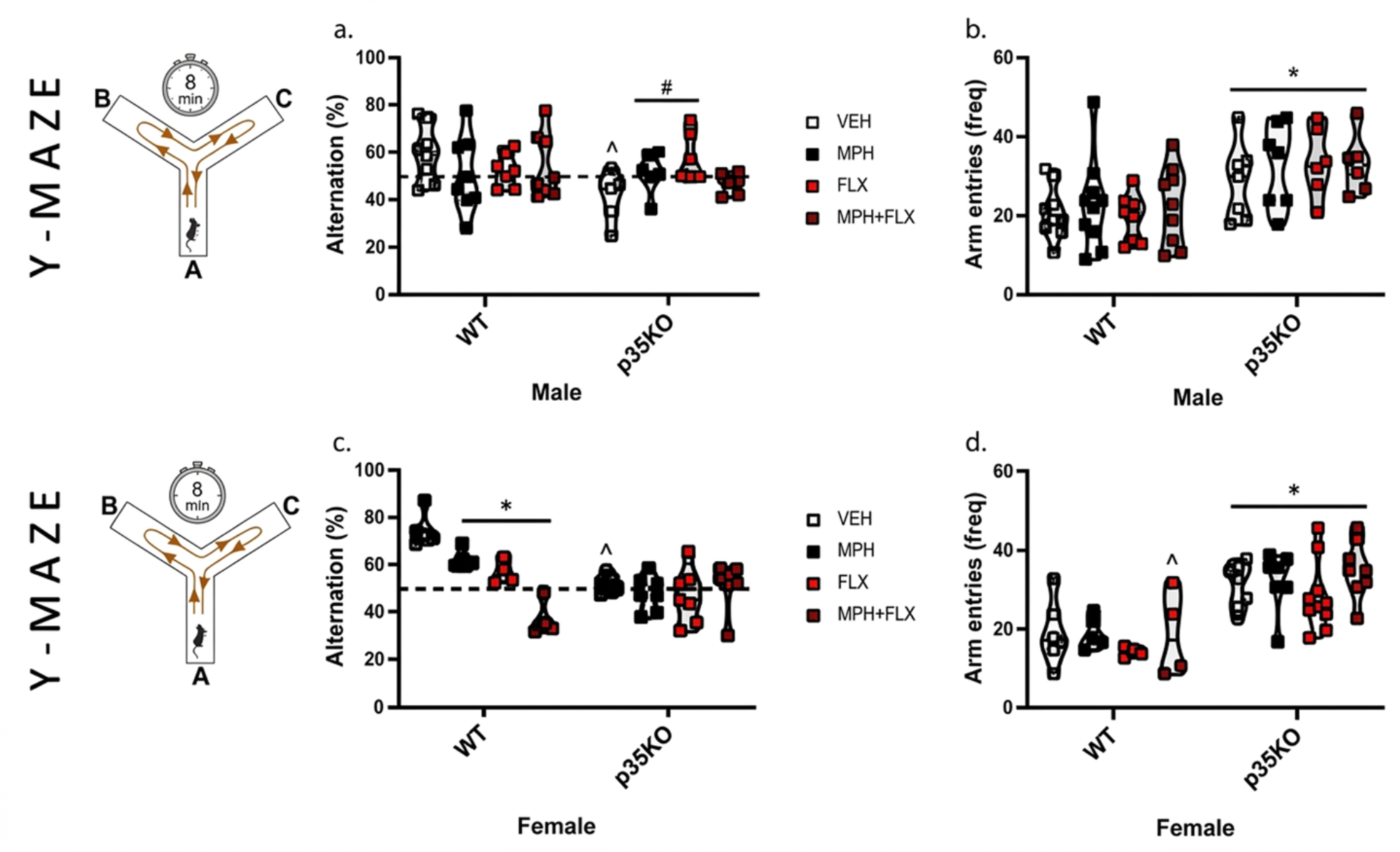
Effects of psychoactive drug treatments on working memory performance in the Y-maze. **Males:** (a) Alternation percentage: ^p < 0.001 (p35KO-VEH vs. WT-VEH); &p < 0.01 (p35KO-MPH and p35KO-FLX vs. p35KO-VEH). (b) Arm entry frequency: *p < 0.05 (p35KO vs. WT). **Females**: (c) Alternation percentage: ^p < 0.001 (p35KO-VEH vs. WT-VEH); £p < 0.01 (WT-MPH, FLX and MPH+FLX vs. WT-VEH). (d) Arm entry frequency: *p < 0.05 (p35KO vs. WT). Data are expressed as mean ± SEM. Statistical analyses were performed using three-way ANOVA followed by Tukey’s post hoc test.

Pharmacological treatment modulated performance in a genotype-dependent manner. In p35KO mice, administration of MPH or FLX significantly increased spontaneous alternation relative to vehicle-treated p35KO animals, consistent with improved WM performance (Fig. 5 a, p < 0.01). In contrast, the combined treatment (MPH+FLX) failed to enhance performance, with alternation scores remaining comparable to vehicle-treated p35KO mice.

Conversely, WT mice exhibited significantly reduced alternation following MPH, FLX, or combined treatment relative to vehicle-treated WT controls. This reduction was visually more pronounced in healthy females, although no significant interaction involving sex was detected (Fig. 5 c, p < 0.01).

Exploratory activity, measured as the number of arm entries, was significantly higher in p35KO mice than in WT animals (F(1,54) = 15.500, p = 0.00025), independently of sex or treatment, with no significant genotype × treatment interaction. This increase was consistent across experimental groups and was not modified by pharmacological intervention (Fig. 5 b, d).

## DISCUSSION

Attention-Deficit/Hyperactivity Disorder (ADHD) is characterized by a persistent pattern of inattention, hyperactivity, and impulsivity that interferes with academic performance, social functioning, and emotional regulation. These behavioral symptoms arise from complex neurobiological mechanisms underlying the cognitive deficits associated with the disorder. Among the molecular pathways implicated, Cdk5/p35 signaling plays a pivotal role in prefrontal cortical function and WM. Here, we show that dysregulation of this pathway in p35KO mice, which exhibit reduced Cdk5 activity (Hallows et al., 2003), leads to altered patterns of c-Fos-IR expression and differential behavioral responses to psychoactive drugs, supporting the use of this animal model with ADHD-like phenotypes to investigate neurobiological processes relevant to cognitive dysfunction associated with ADHD.

ADHD is also strongly influenced by sex, with males diagnosed more frequently than females, showing male-to-female ratios of approximately 3:1 to 5:1 (Martin et al., 2024). This imbalance partly reflects historical biases in diagnostic criteria and clinical assessment, as females often present with subtler or atypical symptoms that are easily overlooked (Mowlem et al., 2019; Nøvik et al., 2006). Despite increasing awareness, women remain less likely to receive pharmacological treatment unless symptoms are severe. Considering these disparities, incorporating sex as a biological variable has become essential for understanding the neurobiological and pharmacological heterogeneity of ADHD. In this context, our study provides experimental evidence highlighting the contribution of sex as a biological variable to behavioral and pharmacological responses using the p35KO mouse model for studying ADHD.

Given the clinical evidence that cognitive symptoms in ADHD vary between sexes, we first examined WM performance in male and female p35KO mice under basal conditions to determine whether the absence of p35 differentially affects this prefrontal-dependent function. p35KO mice showed significant impairment in WM, evidenced by a reduction in spontaneous alternations in the Y-maze test compared to WT counterparts, consistent with the executive dysfunction frequently reported in ADHD patients. These deficits were present in both sexes, indicating that the loss of p35—and consequent reduction in Cdk5 activity (Hallows et al., 2003)—impairs prefrontal-dependent cognitive processing regardless of sex. In contrast, recognition memory, as assessed by the NOR test, remained intact in all groups. Additionally, p35KO mice of both sexes exhibited increased exploratory behavior, a behavioral feature that may reflect enhanced novelty seeking associated with ADHD-like traits. Interestingly, females spent less time exploring novel objects than males, regardless of genotype, underscoring the importance of incorporating sex as a biological variable in ADHD research.

To further explore neural activity associated with the observed behavioral alterations, we analyzed patterns of neuronal activation in prefrontal cortical and hippocampal regions by assessing c-Fos-IR expression as an indirect marker of neuronal activity. Given the central role of these structures in WM and exploratory behavior, this approach allowed us to determine whether the absence of p35 is associated with region- and genotype-dependent patterns of c-Fos-IR expression, with overall effects of sex.

Behavioral testing demonstrated a robust WM impairment in p35KO mice, a cognitive process commonly classified as a “cold” executive function and primarily associated with medial prefrontal cortical processing (Liu et al., 2018; Nordquist et al., 2003; Salehinejad et al., 2021). In this framework, cold executive functions refer to cognitively driven processes such as WM and attentional control, whereas “hot” executive functions involve value-based and motivational aspects of behavior (Salehinejad et al., 2021). The c-Fos-IR analysis revealed that our model exhibits a region- and sex-specific pattern of neuronal activation associated with task performance. In parallel, p35KO animals exhibit altered cortical layering (Chae et al., 1997), consistent with delayed prefrontal neurodevelopment similar to that described in ADHD patients (Shaw et al., 2007), suggesting early alterations in the organization of cortical circuits supporting cognitive control. Within this conceptual framework, the observed patterns of prefrontal activation are consistent with differential engagement of prefrontal subregions in relation to the behavioral phenotype.

In p35KO mice, we observed reduced post-task c-Fos-IR expression in prelimbic and infralimbic cortices, regions commonly associated with WM processes involving the maintenance and manipulation of information. We also detected reduced c-Fos-IR expression in the orbitofrontal regions, which have been linked to value-related and motivational aspects of executive function. This pattern is broadly consistent with frameworks that consider these areas important for controlled cognition and goal-directed behavior (Friedman and Robbins, 2022).

In contrast to these prefrontal patterns, the anterior cingulate cortex (Cg1)—often described as integrating cognitive and motivational aspects of executive function and contributing to conflict monitoring and performance evaluation—showed comparable activation levels in p35KO and WT mice of both sexes. This preserved activation indicates that some aspects of task-related engagement remain detectable in this region. Together, these findings suggest that p35 deletion is associated with more prominent changes in specific prefrontal areas involved in ‘cold’ executive functions associated with WM than in the Cg1.

The p35KO mice, also exhibited increased hippocampal c-Fos-IR in CA1, CA3, and dentate gyrus alongside reduced activation in prefrontal regions. This pattern indicates a differential engagement of hippocampal and prefrontal areas during task performance. Such regional differences in task-related neuronal activity, inferred from c-Fos-IR expression, may be associated with the WM deficits observed in p35KO.

Although no significant interactions involving sex were detected, some differences between males and females were observed in specific conditions. These differences may reflect compensatory mechanisms or variations in baseline activity in WT animals. Importantly, given that prior studies using the p35KO model have been conducted predominantly in males, our findings extend its characterization by revealing marked differences related to sex, in both behavioral and pharmacological responses. This interpretation is consistent with previous evidence of sex dimorphism in the functional organization of prefrontal networks, highlighting the importance of systematically incorporating both sexes in future studies using this model. Large-scale neuroimaging studies have demonstrated that intrinsic brain networks exhibit consistent sexual dimorphism, particularly within task-positive control networks that underlie cognitive control and attentional processes (de Lacy et al., 2019). At a more local level, females show enhanced glutamatergic transmission in the mPFC—reflected in higher amplitude and frequency of spontaneous excitatory postsynaptic currents and greater inward rectification—which suggests a stronger excitatory drive that may buffer against reductions in prefrontal activation (Knouse et al., 2022). Additionally, sex differences in intracortical network activity have been observed ex vivo from prepubertal stages through aging, indicating stable and developmentally conserved divergence in prefrontal network dynamics (Sigalas et al., 2017).

In summary, p35 deletion is associated with region-specific differences in neuronal activation, characterized by reduced prefrontal and increased hippocampal activation in males, while females showed a distinct activation profile. These findings support the use of the p35KO model to investigate neurobiological processes relevant to cognitive dysfunction associated with ADHD, with differences related to sex. In this context, WM relies on coordinated activity across these regions (Liu et al., 2018; Miller and Cohen, 2001; Tamura et al., 2017), and the altered activation patterns observed in p35KO mice are consistent with network-level differences accompanying the cognitive phenotype.

The pharmacological findings reveal genotype-dependent differences in behavioral responses to psychoactive drugs in the p35KO model, and they also suggest differences in drug responses across groups, including effects related to sex. Improvements in WM in p35KO were observed only in male mice treated with MPH or FLX alone, whereas combined treatment did not produce similar effects. These observations indicate that treatment outcomes differ across biological groups rather than producing uniform cognitive changes.

This improvement parallels the “paradoxical response” to psychostimulants previously described in male mice of this model, where drugs such as MPH or amphetamine reduce hyperactivity and normalize dopaminergic signaling, in contrast to their activating effects in WT animals (Krapacher et al., 2010). Previous evidence indicates that Cdk5/p35 dysfunction alters dopamine homeostasis by increasing striatal dopamine content while reducing dopamine turnover and surface DAT expression (Fernández et al., 2021; Krapacher et al., 2010). Thus, the beneficial effects of MPH on WM may reflect a partial restoration of dopaminergic tone within fronto-striatal circuits (Berridge et al., 2006), a mechanism that could also contribute to the observed effects of FLX through serotonergic modulation of prefrontal and hippocampal pathways.

The absence of improvement with the combined MPH+FLX treatment is particularly noteworthy. Several studies have shown that co-administration of SSRIs with psychostimulants can potentiate gene regulation in striatal regions, increase stereotyped behaviors, and facilitate cocaine self-administration (Lamoureux et al., 2024, 2023; Steiner et al., 2010). Such molecular and behavioral potentiation may counteract the therapeutic benefits of each drug when given alone, possibly through excessive activation of overlapping monoaminergic pathways (dopaminergic and serotonergic) or by promoting maladaptive plasticity within reward-related networks. In this context, the lack of cognitive improvement observed in male p35KO mice after combined treatment may reflect altered interactions between dopaminergic and serotonergic systems, consistent with these preclinical findings.

Together, these findings emphasize that sex and neurobiological variability critically shape responses to psychoactive drugs, and caution against assuming that drug combinations effective in one experimental or clinical context will yield similar benefits across sexes or within the heterogeneous ADHD population.

An important finding emerged from the evaluation of WT, which showed a pronounced decline in WM after treatment with MPH, FLX, or their combination, and this effect was particularly evident in females; revealing possible differential pharmacological sensitivity related to sex in healthy animals. Previous reports have emphasized that the effects of MPH differ markedly between control and ADHD models, which is crucial for distinguishing its therapeutic versus misuse/abuse consequences. For instance, Coelho-Santos et al. (2018) demonstrated that MPH misuse negatively affects rat brain function and behavior (Coelho-Santos et al., 2018). Furthermore, these authors reported that healthy animals exposed to high doses of MPH, support a non-linear, inverted-U relationship between catecholaminergic activity and cognitive performance (Coelho-Santos et al., 2019). In such cases, supra-optimal doses elevate dopamine and norepinephrine levels beyond the range necessary for optimal prefrontal functioning, leading to cognitive disruption. The acute dosing used here may have exceeded the optimal threshold in WT females, resulting in impaired WM performance. Moreover, sex differences in monoaminergic regulation and receptor expression could further modulate drug sensitivity, as previously shown in both rodents and humans (Bangasser and Cuarenta, 2021; Becker et al., 2017).

Finally, the finding that none of the drug treatments altered the increased exploratory behavior of p35KO mice indicates that the pharmacological effects observed were specific to cognitive domains.

Together, these findings indicate that disruption of Cdk5/p35 signaling is associated with WM deficits, region-specific patterns of neuronal activation, and potential variability in pharmacological responsiveness associated with sex. In particular, p35 deletion was linked to reduced prefrontal and increased hippocampal activation, along with distinct behavioral responses to psychoactive drugs across biological groups. The observation that identical pharmacological treatments yielded different outcomes depending on genotype, and possibly sex, underscores the importance of baseline neurobiological context when evaluating treatment effects.

Overall, these findings support the use of the p35KO model as a framework to examine variability in cognitive performance and pharmacological responses related to sex, and highlight the importance of considering biological sex and neurobiological context in preclinical and translational ADHD research, as well as for reconsidering combination therapies that simultaneously target multiple monoaminergic systems.

## ACKNOWLEDGEMENTS

We would like to thank **María Julia Cambiasso** for her invaluable and all-encompassing support, as well as **Claudia Bregonzio** and **Fabiola Macchione** for generously providing the antibodies used in this study. We are especially grateful to **Patricio Pereyra** for his assistance with animal husbandry, and to the vivarium technicians —**Romina Maiorano, Milagros Nigro, Jesica Piovano, Joaquin Nigro and Eliana Martinez**— for their daily support. We also thank **Evelin Cotella** and **Mercedes Benedetto** for their helpful discussions. Technical assistance with the PhenoImager™ Fusion microscope (Akoya Biosciences) was provided by **Soledad Miró** at the Centro de Micro y Nanoscopía de Córdoba, CEMINCO-CONICET-Universidad Nacional de Córdoba, Córdoba, Argentina.

## DATA AVAILABILITY

Datasets generated during the current study are available from the corresponding author upon reasonable request.

## AUTHOR CONTRIBUTIONS

MGP, AG and SL conceived and designed the study. FD and OMB performed the behavioral and pharmacological experiments, conducted c-Fos immunohistochemistry, and conducted data curation and statistical analyses. DYS and MES conducted the NOR experiments. GB quantified c-Fos immunohistochemistry images and FAH analyzed behavioral videos. Supervision and project administration were done by MGP and funding’s acquisition by MGP. JV, AG, FD and MGP interpreted the data and drafted the manuscript. All authors critically revised the manuscript, approved the final version, and agree to be accountable for all aspects of the work.

## DECLARATION OF GENERATIVE AI AND AI-ASSISTED TECHNOLOGIES IN THE MANUSCRIPT PREPARATION PROCESS

During the preparation of this manuscript, the authors used ChatGPT (OpenAI) to improve language and readability. The authors reviewed and edited the content and take full responsibility for the content of the publication.

## Supplementary Material and Methods

### c-Fos-IR immunohistochemistry

c-Fos-IR expression was used as an indirect marker of task-related neuronal activity and was assessed by immunohistochemistry, following previously described procedures [25]. Briefly, WT and p35KO mice were anesthetized by i.p. injection with an overdose of 30% chloral hydrate (0.1 mL/100 g) and transcardially perfused with isotonic saline solution followed by 4% paraformaldehyde in 0.1 M phosphate buffer, pH 7.2 (Riedel-de Haën, Sigma-Aldrich Laborchemikalien GmbH, Seelze, Germany). The brains were then removed, fixed overnight in the fixative solution, and stored at 4°C in 30% sucrose until processing and coronal sections of 40 µm were cut using a freezing microtome (Reicher-Jung Hn40, Leica Microsystems, Wetzlar, Germany). The Fos antibody used in this study was raised in rabbits against a synthetic 14-amino acid sequence corresponding to residues 4–17 of human Fos (Ab-5, batch no. 60950101; Oncogene Science, Manhasset, NY). Following primary antibody incubation, sections were washed three times in PBS and placed in 1:200 goat anti-rabbit IgG biotinylated antibody (Vector Laboratories, Burlingame, CA, USA) in 1% NHS PBS for 2 h. The sections were then washed 3 times in PBS and incubated in avidin-biotin complex (Vectastain, ABC Kit, Vector Laboratories) for 2 h. After washing 3 times in PBS, sections were immersed in 0.05% 3,3’-diaminobenzidine tetrahydrochloride and 0.005% H2O2 to visualize the reaction product. The reaction was stopped with PBS, and sections were mounted on subbed slides, dried on a slide warmer, dehydrated and coverslipped with DPX (Fluka, Biochemika, Sigma Aldrich, Chemie GmbH, Steinheim, Germany).

### Anatomical Context and Region Selection

Coronal sections were analyzed at four distinct anteroposterior (AP) levels relative to Bregma, each corresponding to a predefined analysis category within the macro (Table 1). The “Analyze Image” function automatically determined the number of ROIs for quantification based on the user’s initial input specifying the anatomical level of the section.

**Table 1.** Anatomical regions and corresponding ROI count defined by the macro based on the selected anteroposterior level.

| AP Level<br>(Bregma) | Macro Label<br>(Internal Name) | N<sub>ROIs</sub> | Target Regions Analyzed |
| --- | --- | --- | --- |
| +2.22 mm | Cortex | 5 | Cingulate Cortex (Cg), Prelimbic Cortex (PL), Infralimbic Cortex (IL), Lateral Orbital Cortex (LO), Ventral Orbital Cortex (VLO) |
| -1.58 mm | Hippocampus | 3 | CA1, CA3, Dentate Gyrus (DG) |

### Image Preprocessing and Quantification

Images were acquired using an Akoya Biosciences Phenoimager Fusion equipped with a UPlanXApo 10×/0.40 objective (OFN 26.5). Selected brain regions were preprocessed using the *image preprocess* function, which automated the generation of a binary mask of c-Fos–immunoreactive (c-Fos-IR) nuclei while incorporating user-assisted quality control procedures. To prepare the images, the original files were duplicated and subjected to a Bandpass Filter (large = 40, small = 3) followed by Unsharp Masking (radius = 3, mask = 0.70) to enhance nuclear signal intensity and suppress low-frequency background variation. A semi-automated thresholding step was then applied using the Intermodes algorithm, allowing manual adjustment of the threshold prior to conversion of the image into a binary mask.

The resulting mask was refined through standard morphological image analysis to accurately delineate individual nuclei. Morphological opening was first performed to separate aggregated particles, followed by size filtering (20–500 pixels), Watershed segmentation, and hole filling to complete nuclear contours. A final particle-filtering step retained only objects within a defined size range (30–500 pixels) and shape range (circularity = 0.80–1.00). Manual quality control (QC) was conducted in two phases: during the nuclei-inclusion phase, the experimenter could manually add missed particles using a 15-pixel white pencil, whereas during the artifact exclusion phase, incorrect or spurious objects could be removed using a 15-pixel black pencil.

Region-specific quantification was subsequently performed using the *analyze image* function. The corresponding region of interest (ROI) file was first loaded, allowing manual adjustment of the polygonal ROI boundaries to ensure precise anatomical alignment. Thereafter, both the occupied area and the number of c-Fos-IR nuclei were automatically measured for each ROI. Finally, the procedure generated two CSV output files—one containing the nuclei count and size data, and another containing the total ROI areas—along with a ZIP archive containing the processed image and the final set of ROIs used for quantification.

The ImageJ macro used for preprocessing and quantification was validated in representative sections by comparing automated and manual counts performed by an independent blinded observer, ensuring accuracy and reproducibility, and is available upon request.

**Figure S1.**
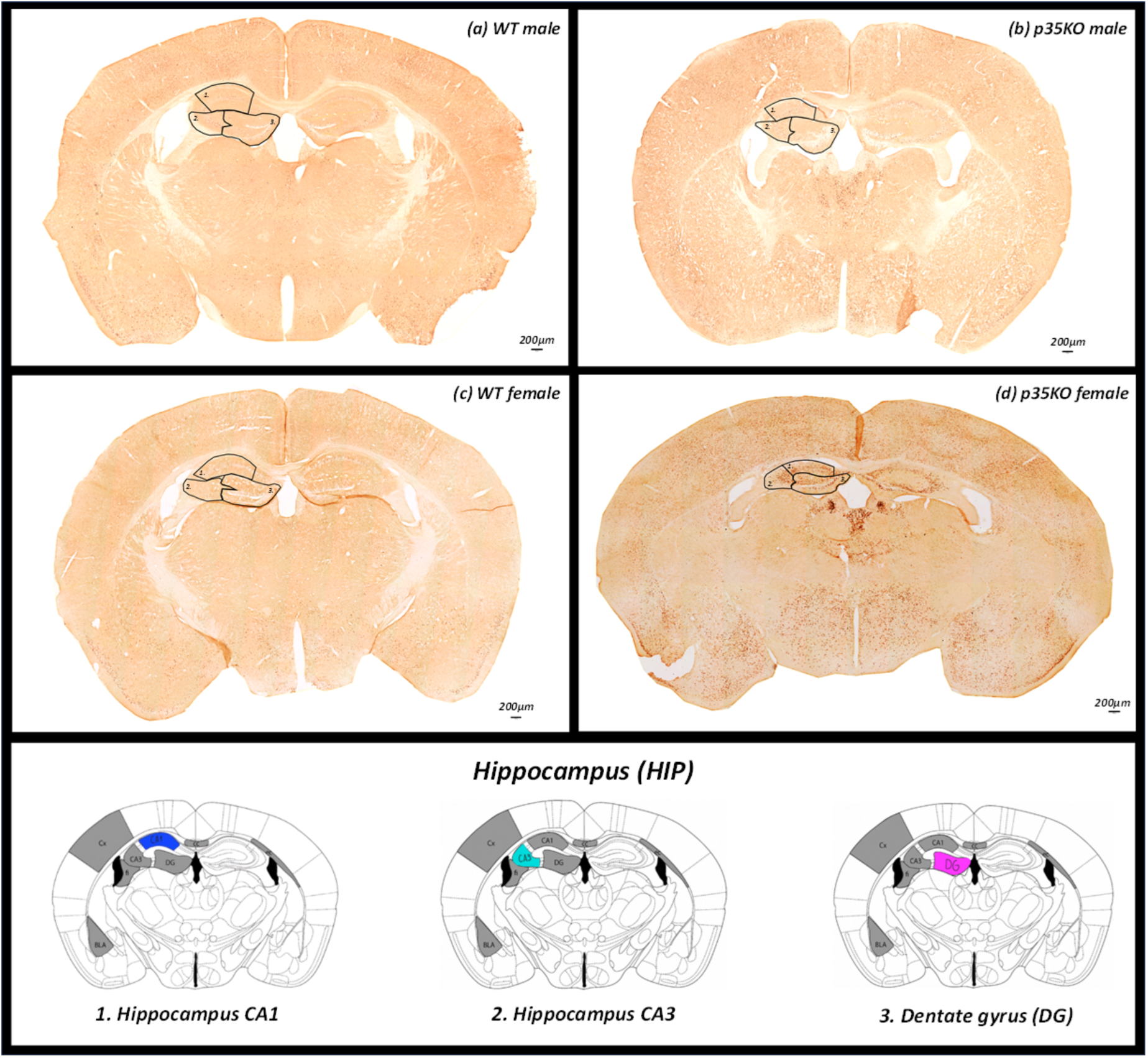
Representative c-Fos-IR immunostaining in the hippocampus of WT and p35KO mice (males and females). Representative photomicrographs show c-Fos-IR expression patterns across hippocampal subregions (CA1, CA3, and dentate gyrus, DG) in WT and p35KO mice of both sexes, obtained 90 min after Y-maze testing. Panels display c-Fos–immunoreactive nuclei in: *(a) WT male, (b) p35KO male, (c) WT female, and (d) p35KO female.* These images provide a qualitative comparison of regional activation across genotypes and sexes. All photomicrographs were acquired using identical acquisition parameters to ensure cross-sample comparability. Olympus UPlanXApo 10x/0.40∞/0,17/OFN 26.5. Scale bar: 200 μm.

**Figure S2.**
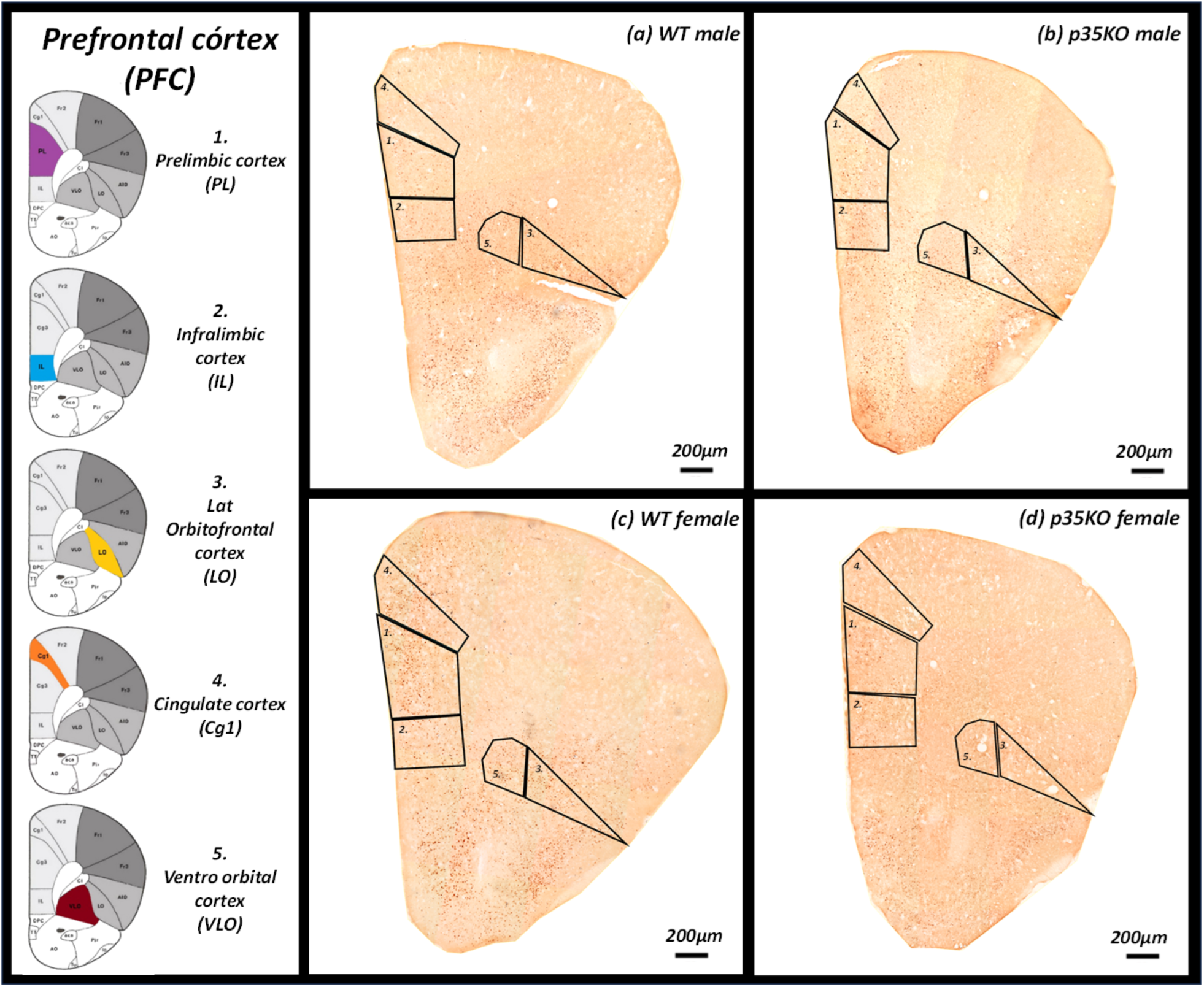
Representative c-Fos-IR immunostaining in the prefrontal cortex of WT and p35KO mice (males and females). Representative photomicrographs show c-Fos-IR expression across prefrontal cortex subregions, including the prelimbic (PL), infralimbic (IL), lateral orbitofrontal (LO), cingulate (Cg1), and ventrolateral orbitofrontal (VLO) cortices, in WT and p35KO mice of both sexes, obtained 90 min after Y-maze testing. Panels display c-Fos–immunoreactive nuclei in: *(a) WT male, (b) p35KO male, (c) WT female, and (d) p35KO female*. Images allow qualitative comparison of activation patterns across genotypes and sexes. All photomicrographs were acquired under identical microscope settings to ensure comparability. Olympus UPlanXApo 10x/0.40∞/0,17/OFN 26.5. Scale bar: 200 μm.

## Highlights

1. Mice lacking p35 exhibit working memory deficits but intact recognition memory.
2. Prefrontal c-Fos decreases while hippocampal expression increases in p35KO mice.
3. MPH or FLX improve working memory only in p35KO males.
4. Combined MPH+FLX treatment abolishes drug-induced improvement in p35KO mice.
5. WT females show marked working memory impairment after drug treatment.

